# Crosstalk between chromatin and the transcription factor Shavenbaby defines transcriptional output along the *Drosophila* intestinal stem cell lineage

**DOI:** 10.1101/2023.01.03.522588

**Authors:** Alexandra Mancheno-Ferris, Clément Immarigeon, Alexia Rivero, David Depierre, Nicolas Chanard, Olivier Fosseprez, Gabriel Aughey, Priscilla Lhoumaud, Julien Anglade, Tony Southall, Serge Plaza, Olivier Cuvier, François Payre, Cédric Polesello

**Author notes:** correspondence (O.C.), (C.P.). these authors contributed equally to this work (Alexandra Mancheno-Ferris, Clément Immarigeon).

## Abstract

The transcription factor Shavenbaby (Svb), the only member of the OvoL family in *Drosophila*, controls intestinal stem cell differentiation. Post-translational modification of Svb produces two protein isoforms, Svb-ACT and Svb-REP, which promote intestinal stem cell renewal or differentiation, respectively. Using engineered cell lines, we express either isoform to define their mode of action, and develop an unbiased method to identify Svb target genes in intestinal cells. Within a given cell type, Svb-ACT and Svb-REP antagonistically regulate the expression of a set of target genes, binding specific enhancers whose accessibility is constrained by. During intestinal differentiation, the set of target genes progressively changes, together with chromatin accessibility. Moreover, Svb-REP binding stabilizes three-dimensional enhancer-promoter loops, while influencing the local chromatin landscape to repress target genes. We propose that SvbACT-to-REP switch promotes enterocyte differentiation of intestinal stem cells through direct gene regulation and chromatin remodeling.

## Introduction

Transcription factors (TFs) are primary determinants of cell phenotypes and behaviors, regulating gene expression via binding to DNA sequences located within enhancers and promoters cis-regulatory elements (Takahashi and Yamanaka, 2006) (Bulyk, 2003; Rohs *et al*., 2009). TF binding motifs being generally short (4-15 base pairs) and degenerate, they are very abundant in the genome. The accessibility of TF binding sites is influenced by many parameters and dimensions, including biophysical constraints, chromatin state and organization into domains, and notably involving the presence/absence of epigenetic co-regulators. As a consequence, only a limited subset of TF binding sites are bound in a given cell type (Buenrostro *et al*., 2018; Cuvier and Fierz, 2017)(Guertin and Lis, 2010; Slattery *et al*., 2014; Wang *et al*., 2012), and substantial variations in binding events are often observed across tissues (Arvey *et al*., 2012; Spitz and Furlong, 2012). These cell-specific changes in binding site occupancy are often thought to have strong impact in transcriptional outputs (Sen *et al*., 2019), though distinct tissues, developmental stages and/or transcriptional factors may adopt distinct strategies. The underlying mechanisms and functional consequences currently remain to be fully elucidated.

Transcription factors of the Ovo-like (OvoL) family are specific to metazoans (Kumar *et al*., 2012) and display key functions during animal development. The founding member called Ovo/Shavenbaby (Svb) was initially identified in flies (Mevelninio *et al*., 1995) for its dual role in germline and epidermal development. Mammalian species have evolved three paralogs referred to as OvoL1-3 that display partly redundant functions (Teng *et al*., 2007) and also contribute to germline, skin development and diseases (Tsuji *et al*., 2017). In addition, the deregulation of OvoL expression is associated to various cancers of epithelial origin (Roca *et al*., 2013; Wang *et al*., 2017; Watanabe *et al*., 2014) and it has been proposed that OvoL factors act as the guardians of epithelial integrity by preventing epithelial-mesenchymal transition (Hong *et al*., 2015; Wang *et al*., 2017). While OvoL TFs share a highly-conserved zinc finger DNA binding domain that recognizes the core sequence CNGTT (Lee and Garfinkel, 2000; Lu and Oliver, 2001), they differ in their N-terminal regions and can exhibit distinct transcriptional activities (Watanabe *et al*., 2014).

*Drosophila* encodes a unique Ovo/Svb gene, providing an attractive paradigm to dissect OvoL functions and investigate the mechanisms underlying tissue-specific activities. The somatic factor, Shavenbaby (Svb), is known for governing differentiation of epidermal cells in the embryo and adult derivatives (Chanut-Delalande *et al*., 2014; Payre *et al*., 1999). The Svb protein is translated as a large transcriptional repressor, referred to as Svb-REP. Remarkably, under the action of Polished-rice (Pri, a.k.a. Tarsal-less or Mille patte) peptides encoded by small open reading frames, the ubiquitin ligase Ubr3 binds to Svb-REP, and triggers proteasome-dependent processing resulting in the production of a shorter transcriptional activator, Svb-ACT (Kondo *et al*., 2010; Zanet *et al*., 2015). In the embryonic epidermis, Svb-ACT triggers the expression of a battery of effector genes that are collectively responsible for the remodeling of epithelial cells (Chanut-Delalande *et al*., 2006; Fernandes *et al*., 2010; Menoret *et al*., 2013). We have recently reported that Svb is critically required for the maintenance of adult epithelial stem cells, which ensure the homeostasis of the renal (Bohère *et al*., 2018) and intestinal systems (Al Hayek *et al*., 2021). In addition, the Svb transcriptional switch operated by Pri peptides is also at work to control the behavior of adult stem cells : SvbACT promotes intestinal stem cell self-renewal and proliferation, while in contrast SvbREP dictates differentiation into enterocytes (Al Hayek *et al*., 2021).

Here we use genomic and bioinformatic approaches to study the molecular action of the two Svb isoforms on gene expression, across the intestinal stem cell lineage. Using engineered cells in culture, we define the molecular mode of action of Svb-ACT and REP. We show that, constrained by chromatin landscape, both Svb-ACT and Svb-REP bind to a same array of open enhancers, mediating antagonistic effects on the expression of a common set of target genes. We develop unsupervised analyses of differential gene expression in response to Svb-ACT and Svb-REP allowing the identification of Svb target genes in adult intestinal stem cells (ISCs) and their progeny enteroblasts and enterocytes. Strikingly, Svb target genes are different in each cell type of the lineage. Interestingly, this change in the target genes repertoire during differentiation is accompanied by chromatin accessibility remodeling, and concomitant accumulation of SvbREP. We further show in cultured cells that Svb-REP participates in remodeling local chromatin landscape, playing an active role in transcriptional repression, which can occur through distal enhancers. This work sheds light on the possible crosstalk between the chromatin landscape co-evolving with the proteasome mediated maturation of a transcription factor, which may serve to optimize transcriptional output to differentiation.

## Results

### Svb-REP and Svb-ACT bind common enhancers and antagonistically control gene expression

To map the genome-wide set of sites bound by Svb-ACT and Svb-REP, we made use of stable cell lines derived from cultured S2 *Drosophila* cells, since they do not express endogenous Svb and have been engineered to express one or the other Svb form (Kondo *et al*., 2010; Zanet *et al*., 2015). We performed chromatin immunoprecipitation coupled to next-generation sequencing (ChIP-seq) in Svb-ACT::GFP or Svb-REP::GFP cells (Figure 1A). ChIP-seq revealed 6,939 regions highly enriched for Svb-ACT binding (Figure 1B). Consistent with a binding mediated by the zinc fingers domain, these regions feature DNA motifs matching CcGTT, the Ovo/Svb binding matrixes ((Castro-Mondragon *et al*., 2017; Menoret *et al*., 2013; Nguyen *et al*., 2018) and methods). The most enriched motif matches a ACCGTTA sequence (hereafter referred to as F7-BS, Figure 1D). F7-BS - defined by a machine learning approach combining statistical analysis to phylogenetic information (Rouault *et al*., 2010) is the best predictor of functional Svb binding sites *in vivo* (Menoret *et al*., 2013).

**Figure 1:**
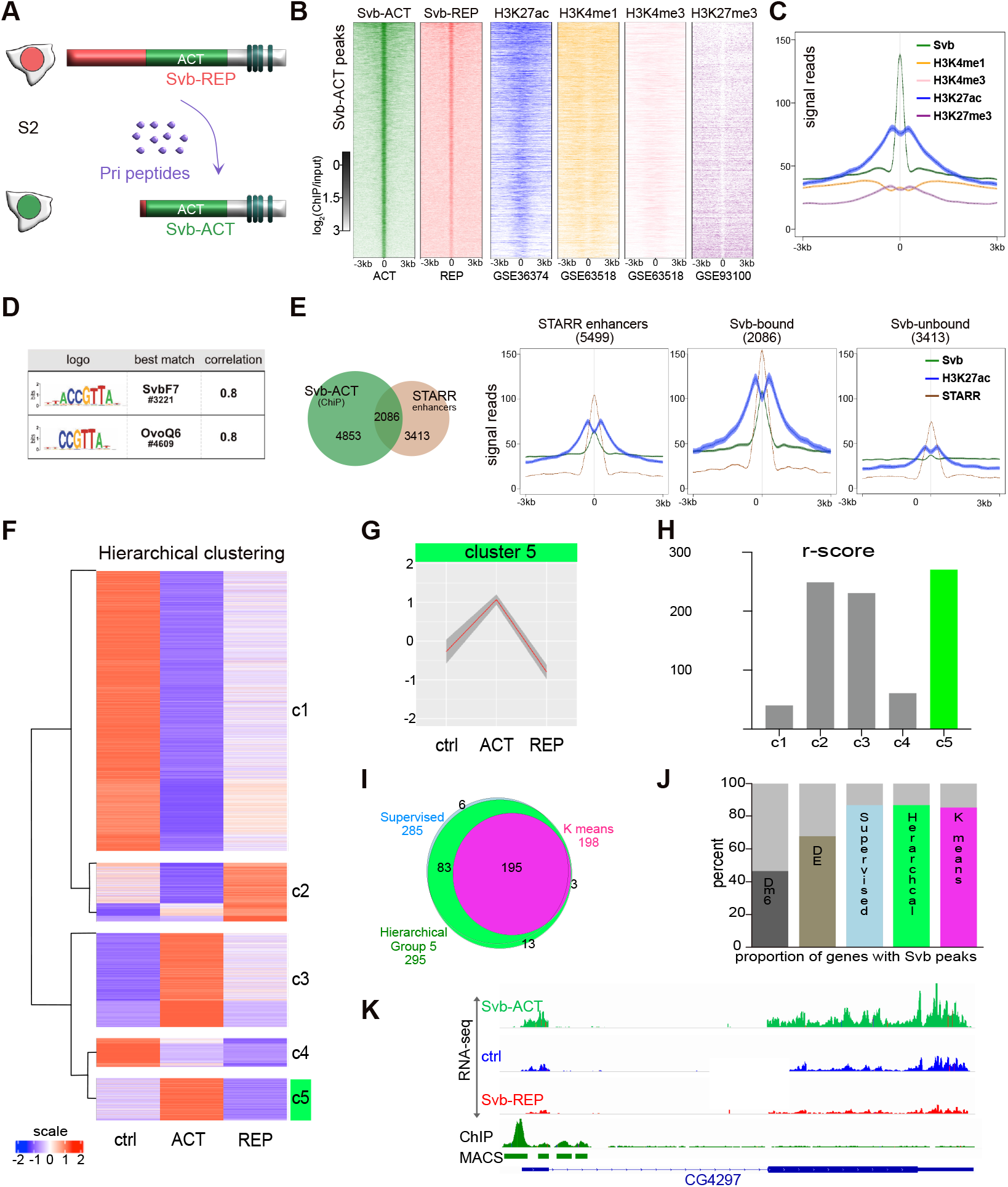
Svb-ACT and REP bind common enhancers to antagonistically regulate a common set of target genes. (A) Cultured S2 cells provide an exquisite model system for studying the mechanistics of Svb isoforms functions. Pri smORF peptides trigger SvbREP to SvbACT post-translational, proteasome-dependent processing. (B) ChIP-seq heatmaps for Svb-ACT, SvbREP and the indicated histones marks available from S2 cells. All heatmaps are centered on Svb-ACT peaks and sorted by Svb-ACT signal intensity. Color intensity reflects the level of ChIP-seq enrichment presented as log2(ChIP/input). (C) Histone marks averaged profile centered on Svb-ACT peaks. (D) DNA motifs most enriched in Svb-ACT bound peaks as defined by RSAT suite, the correlation score reflect the result of the Pearson test between the logo detected and the logo present on the RSAT database (Nguyen *et al*., 2018). (E) Interaction between Svb-ACT peaks and active enhancers identified by STARR-seq (Arnold *et al*., 2013). Left, venn diagram showing the overlap of Svb-ACT bound regions (green, n=6939) and active enhancers (brown, n=5499). Right, average enrichment of Svb-ACT (green), STARR signal (brown) and H3K27ac (blue) for all active enhancers, Svb-bound and Svb-unbound enhancers. (F) Hierarchical clustering of DE genes across RNAseq experimental conditions. (G) Metaprolife of the Cluster 5 of the hierarchical clustering identified in F. (H) r-score defines the robustness of each gene cluster from hierarchical clustering, based on its conservation through 15 iterations of k-means clustering. See Figure S1 and methods for details. c5 is highlighted in green. (I) Venn diagram showing a robust set of 195 genes considered as “S2 Svb target genes” is found through supervised (see Figure S1), hierarchical and k-means clusterings. (J) Diagram showing the proportion of genes with SvbACT ChIP peaks in the whole genome (Dm6), all differentially expressed genes (DE), or Svb putative target genes define by either supervised method, hierarchical clustering or K-means clusterings. (K) Genome browser screenshot of the CG4297 locus displaying typical Svb target gene behavior (basal expression in blue, Svb-ACT upregulation in green; Svb-REP downregulation in red, Svb ChIP in dark green).

Although peak calling detected a markedly smaller number of regions for Svb-REP (1,325), they extensively overlap with Svb-ACT bound regions and there is clear enrichment for Svb-REP across all Svb-ACT peaks (Figure 1B, Figure S1A-F)). We thus conclude that both isoforms bind common sites genome-wide.

To characterize the chromatin landscape of Svb-bound regions, we interrogated whether these regions display recognizable patterns of histone marks using the large set of data available in *Drosophila* cells (Herz *et al*., 2012; Huang *et al*., 2017; Kellner *et al*., 2012; Li *et al*., 2015; Schauer *et al*., 2017). Considering all Svb-ACT peaks (allowing peak detection at higher sensitivity), a strong enrichment in histone3 acetylated on K27 (H3K27ac) across sites bound by Svb was found (Figure 1B-C). Notably, H3K27ac enrichment directly correlated with Svb-ChIP signal (see the intensity “gradients” in Figure 1B). H3K27ac marks open chromatin, in particular active enhancers (Creyghton *et al*., 2010; Cubenas-Potts *et al*., 2017; Li *et al*., 2015) which are also enriched in histone 3 methylated on K4 (Heintzman *et al*., 2009; Sethi *et al*., 2020). Consistently, Svb-bound sites were consistently enriched in H3K4me1 and H3K4me3 that accumulated on both sides of the peak (Figure 1B-C). Accordingly, H3K27me3 marks of repressive chromatin (Liu *et al*., 2020) were depleted (Figure 1B-C). Therefore, these data suggest that Svb binds active enhancers. We challenged this hypothesis by analyzing Svb binding to all enhancers defined by STARR-seq ((Arnold *et al*., 2013) Figure 1E). This revealed that enhancers associated with high Svb signal display stronger enhancer activity (brown) and higher H3K27ac (blue), compared to Svb unbound enhancers (Figure 1E), confirming that Svb binds active enhancers.

To analyze the respective influence of Svb isoforms on gene expression, we next performed RNA-sequencing in control, Svb-ACT and Svb-REP cells. When compared to controls, both Svb-ACT and Svb-REP significantly changed gene expression (Figure S1G). *In vivo* assays in embryonic epidermal cells have shown the antagonistic activity of Svb-ACT and Svb-REP on individual direct target genes (Chanut-Delalande *et al*., 2006; Fernandes *et al*., 2010; Kondo *et al*., 2010), leading to the upregulation or downregulation of those, respectively. We expected direct target genes of Svb in S2 cells to display a similar behavior. Therefore, we selected all differentially expressed genes (DEG, p-value<0.05) between Svb-ACT and Svb-REP, and focused on those responding to both isoforms (log_2_(FC)>0 in Svb-ACT and log_2_(FC)<0 in Svb-REP conditions, Figure S1G). This supervised approach defined a group of 285 putative target genes. Consistent with the prediction that these genes contain direct targets, 86% of those are associated with Svb binding peaks (Figure 1J). To further tackle the influence of Svb REP and ACT on gene expression, we develop an unsupervised approach based on hierarchical clustering and k-means to identify Svb direct target genes solely based on RNAseq data. Hierarchical clustering (Johnson, 1967) defines 5 groups of DE genes (Figure 1F), one being very similar to the supervised set (278/285 genes in common, Figure 1G,I). We used 15 iterations of k-means clustering to challenge the robustness of each hierarchical cluster gene sets, and developed the “robustness score” method to estimate the proportion of genes that cluster together throughout K-means iterations (r-score, Figure 1H, see methods). The cluster comprising putative Svb targets got the highest r-score and 195 genes were common to all iterations and methods (Figure 1I), 86% of these common genes being associated to Svb binding peaks (Figure 1J). This set of 195 genes is referred hereafter as S2 Svb target genes.

Taken together, these data show that Svb-ACT and Svb-REP display antagonistic transcriptional activities, acting on a common set of approximately 200 genes validated through an unsupervised approach (see Figure 1K for a representative target gene).

### Chromatin landscape defines Svb binding sites

Having defined the set of Svb target genes in S2, we next asked whether it reflected the regulation of genes by Svb in embryonic epidermal cells (Menoret *et al*., 2013), focusing on direct target genes (*i*.*e*., being differentially expressed and displaying a binding peak in embryo). No significant overlap between the two sets of genes was observed (Figure 2A). This suggested that Svb could behave differently depending on the cell type. Indeed, this divergence is also observed when comparing Svb-ChIP peaks between embryonic and S2 cells (Figure 2B).

**Figure 2:**
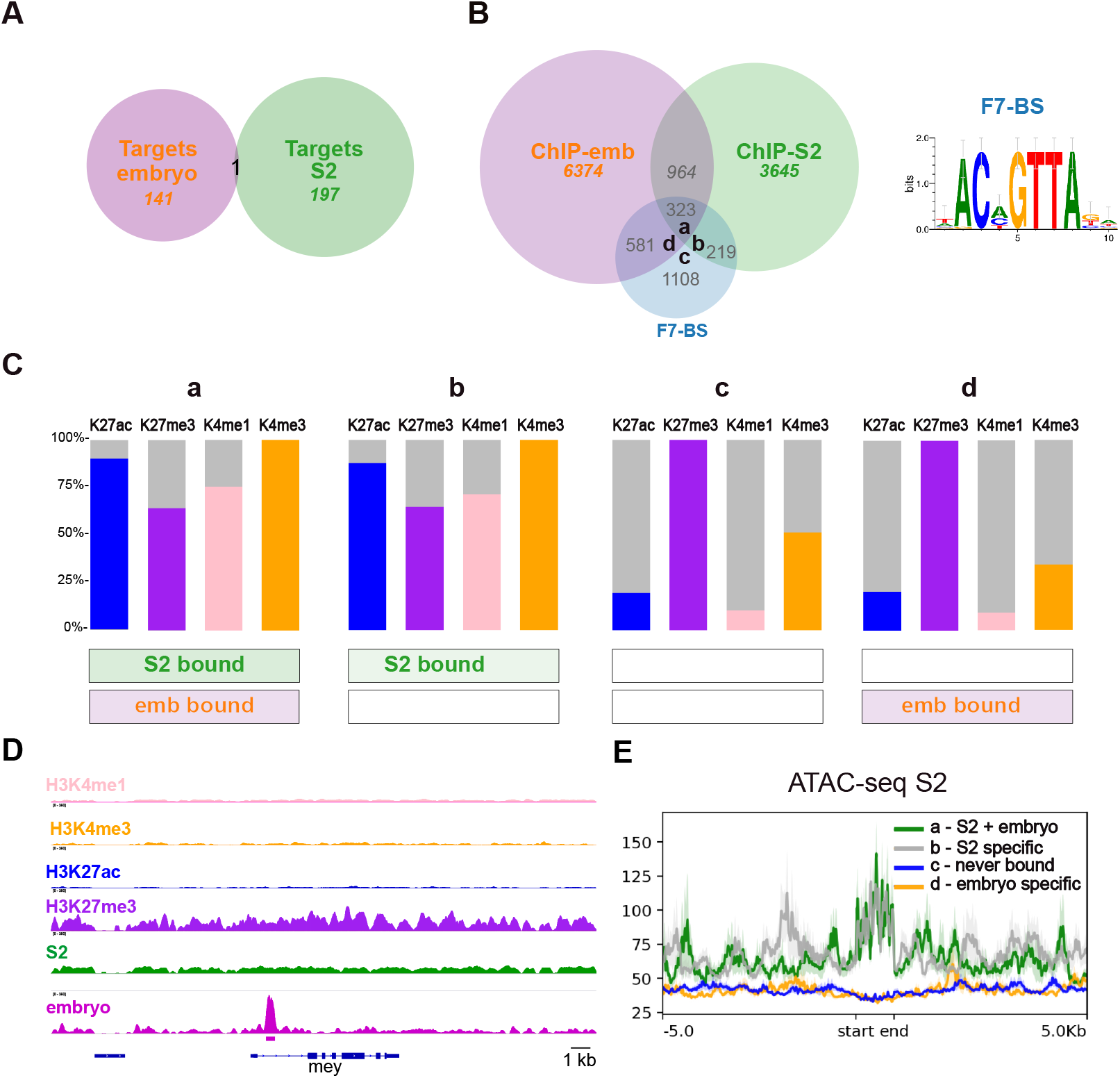
Chromatin landscape constrains Svb binding. (A) Venn diagram showing the overlap between the sets of Svb target genes in S2 cells (n=198) and embryonic epidermal cells (n=142,(Menoret *et al*., 2013)). (B) Venn diagram showing the intersections between Svb-bound regions in S2 cells (green), in embryonic epidermal cells (orange), and genomic F7-BS regions (blue). F7-BS position weight matrix is shown on the right. The “Svb-bound” regions overlap with F7-BS regions if a F7-BS is found less than 3kb away from the ChIP peak. Each F7-BS is thus bound or not by Svb in either cell type. a,b,c and d mark the 4 types of F7-BS behavior shown in panels C and E. (C) Enrichment of the indicated histones marks around S7-BS motifs in the different sets defined in (B). (D) Snapshot of IGV genome browser showing Svb ChIP (from S2 cells in green and from embryos in magenta) at mey locus. In embryo Svb binds a canonical enhancer driving *mey* gene expression in embryonic epidermal cells (Menoret *et al*., 2013). The different histone marks (H3K4me1 in pink, H3K4me3 in yellow, H3K27ac in blue, H3K27me3 in purple) allow visualization of the chromatin organization of the region in S2 cells, consistent with repressive configuration and absence of Svb binding in this cell type. (E) ATAC-seq metaprofiles from S2 cells (Ibrahim *et al*., 2018) show chromatin accessibility over 10 kb around F7-BS for the clusters defined in B.

To get a deeper look at the chromatin context of Svb binding, we focused on evolutionarily conserved F7-BS motifs that often mediate transcription of Svb direct targets *in vivo* (Menoret *et al*., 2013) and are highly enriched within Svb peaks in S2 cells (Figure 1D). There are 2354 conserved F7-BS in the *Drosophila* genome, 26% of those being bound by Svb in S2 cells (Figure 2Ba+b). Chromatin profiles of histone post-translational modifications were generated to inspect the chromatin landscape around F7-BS. This analysis showed that Svb-bound F7-BS (a and b) displayed significantly higher H3K27ac, H3K4me1 and H3K4me3 and lower H3K27me3, as compared to non-bound F7-BS (Figure 2C, c and d). Of note, the binding status of F7-BS in embryo has no effect on the chromatin landscape in S2 cells (a vs b and c vs d): whereas F7-BS bound only in S2 cells also displayed the features of active enhancers (b), sites bound by Svb only in the embryonic epidermis exhibited closed chromatin configuration in S2 cells (d). A representative embryonic Svb target gene is shown (Figure 2D, see repressive chromatin landscape and absence of Svb-ChIP peak in S2). We challenge this observation using available ATAC-seq data from S2 cells, and confirm that chromatin is open and accessible at Svb-bound F7-BS (Figure 2E a and b) whereas Svb-unbound sites are in a closed conformation (Figure 2E c and d), again independently of Svb binding in embryo.

Together, these results indicate that the Svb transcription factor regulates different sets of target genes in S2 cells and embryonic epidermal cells. Specific patterns of histone marks highlight how such cell-type specific genes may be gated through accessibility of distinct pool of sites that are accessible to Svb binding, hence defining a subset of cell-type specific target genes regulated by Svb.

### Identification of Svb target genes across the intestinal stem cell lineage

While the absence of endogenous expression of Svb in cultured cells was helpful to address the binding and transcriptional effect of each isoform, the extent to which this experimental model was representative of the choice of target genes *in vivo* remained to be established. We next investigated its activity in adult intestinal stem cells (ISCs) lineage leading to enterocyte (EC) differentiation (Figure 3A). We previously described that Svb-ACT acts in ISC to promote their maintenance and proliferation, while Svb-REP triggers and accompanies differentiation into ECs (Al Hayek *et al*., 2021). Svb-REP accumulates during differentiation (Figure 3A), and is required in mature EC to maintain their epithelial properties (Al Hayek *et al*., 2021). Thus, the switch from Svb-ACT to REP isoforms has a drastic effect on ISC and EC behavior, but the target genes mediating this function remained unknown.

**Figure 3:**
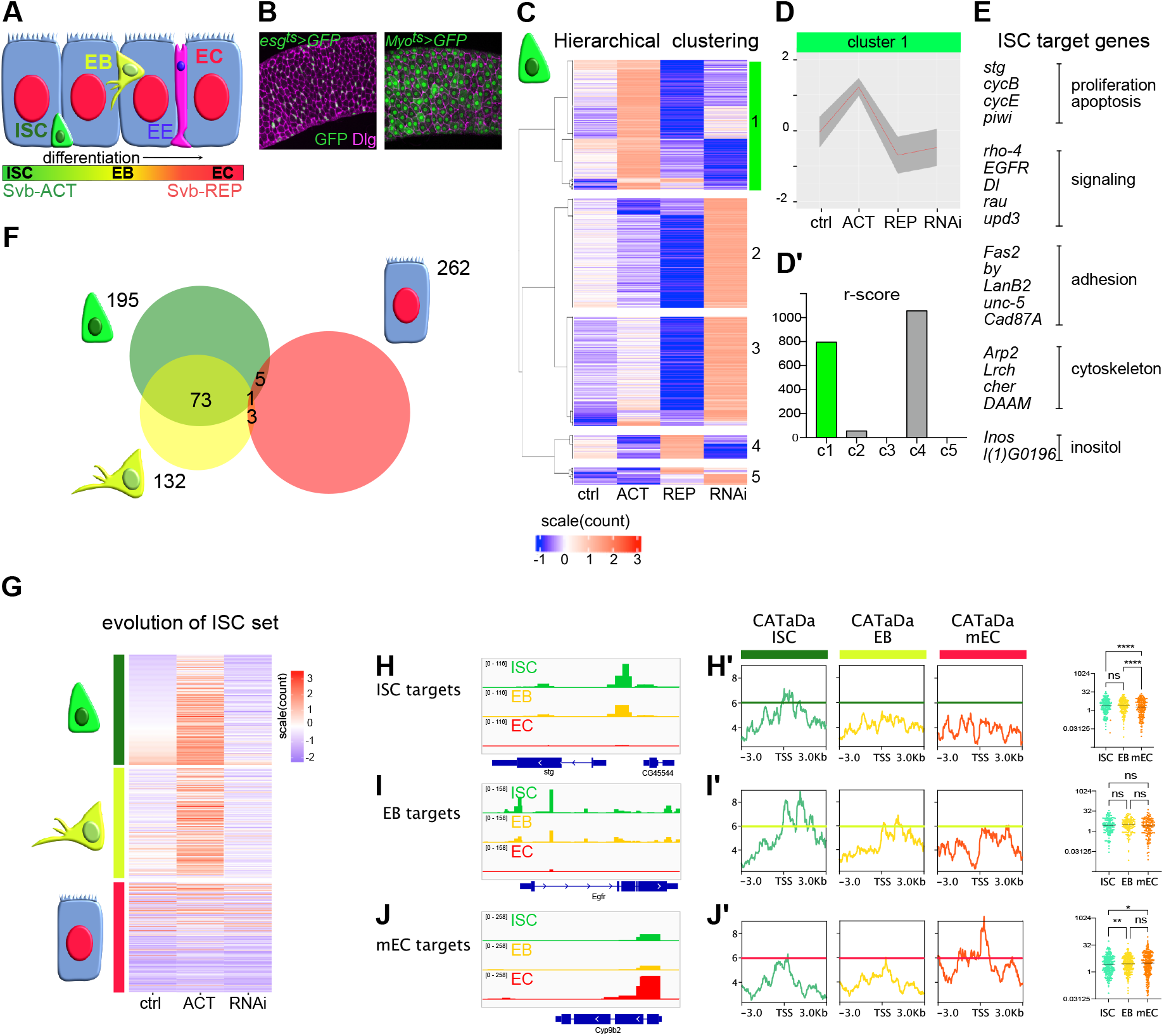
Evolution of Svb target genes through the lineage of intestinal stem cells. (A) Schematic representation of the cell types of the intestinal epithelium leading to EC differentiation. Svb-ACT is the majority form in ISCs and Svb-REP in ECs. ISC in green, EB in yellow, EC in bleue and EE in purple. (B) Cell type specific drivers have been used to manipulate Svb expression in adult fly progenitors (ISCs and EBs, *esg:GAL4, Gal80*^*ts*^) or ECs (*myo1A:GAL4, Gal80*^*ts*^). (C) Hierarchical clustering of RNAseq data from sorted ISC reveals DE genes upon Svb modulation. Four experimental conditions were used: *control* (ctrl), *UAS-SvbACT* (ACT), *UAS-Svb*^*3Kmut*^ (REP), and *UAS-Svb RNAi* (RNAi). Cluster 1 comprises genes activated by SvbACT and repressed by SvbREP and upon Svb loss of function. (D) Metaprofile of c1 corresponding to ISC target genes. (D’) Robustness analysis (r-score) of each hyerarchical cluster. (E) Subset of genes from c1 with their associated functions. (F) Venn diagram showing how Svb target genesets evolve during intestinal cell differentiation. Note ISC/EB target genesets similarity as opposed to EC targets. Svb target genes were identified in ISC, EB, eEC and mEC following the same pipeline (see FigureS2/Table S1). (G) Heatmaps showing expression of the 195 ISC target genes in ctrl, Svb-ACT and *svb-RNAi* conditions, in ISC, EB and mEC (top to bottom). (H-J) CATaDa data (Aughey *et al*., 2018) allow studying the dynamics of chromatin accessibility through the lineage. (H,I,J) are IGV genome browser screenshots of representative loci of an ISC target gene (*stg*, H), an EB target gene (*egfr*, I) and a mEC target gene (*Cyp9b2*, J), showing CaTADA signal in ISC (green), in EB (yellow) and in EC (red). Averaged CATaDa signal profile (in reads per million), centered on the TSS of all ISC target genes (H’), all EB target genes (I’) and all mEC target genes (J’) are shown, showing accessibility in ISC (green), EB (yellow) and EC cells (red) (left to right of each panel). The plots on the right present for each gene the CATaDa signal around the TSS (1kb on both side). Statistical analysis confirms the general trend that ISC target genes’ TSS close and mEC target genes’ TSS open during intestinal cell differentiation (Kruskal-wallis test with Dunn’s multiple comparisons).

To identify genes regulated by Svb in each cell type from the lineage, we profiled the transcriptome of four cell types from posterior midguts: ISCs, enteroblasts (EBs), early EC (eEC) and mature EC (mEC) (see methods). These cells were purified from dissected guts of control animals, upon RNAi-mediated *svb* depletion, and expression of Svb-ACT or Svb-REP^3Kmut^ (a engineered form insensitive to Pri and thus not convertible into Svb-ACT (Zanet *et al*., 2015). Note that manipulation of Svb function was specifically targeted either to progenitor cells (ISC and EB) or to ECs (eEC and mEC), using specific drivers allowing spatial and temporal control (Figure 3B, (Jiang and Edgar, 2009; Micchelli and Perrimon, 2006)).

We first verified that our RNA-seq datasets are consistent with available transcriptomic data and indeed represent distinct cell types along the ISC to EC differentiation route (ISC, EB, early EC and mature EC, Figure S2A). Then, building on the unbiased method developed using cultured cells (Figure 1), we identify the set of genes whose expression is likely directly regulated by Svb. In each cell type, we identify one robust cluster of 132-265 genes antagonistically regulated by Svb as expected for direct target genes (Figure 3C to G, Figure S2B). The fact that their expression was also downregulated upon *svb* knockdown in progenitors confirms that these genes are normally activated by Svb-ACT in ISCs (Figure 3C and G, Figure S2B). On the contrary, Svb target genes in mature ECs tend to be upregulated upon *svb* knockdown, confirming that Svb acts mostly as a repressor in these cells (Figure S2B/Table S1).

In agreement with the function of Svb in ISC (Al Hayek *et al*., 2021), we identify genes controlling cell cycle, proliferation and survival of stem cells (*string, cycE, cycB, piwi*…) as well as signaling pathways known to control stemness (*egfr, Dl*) (Figure 3E) (reviewed in (Boumard and Bardin, 2021)(Kohlmaier *et al*., 2015; Sousa-Victor *et al*., 2017)). This discovery reinforces the importance of the switch between Svb-ACT and REP isoforms in ISC progeny: the downregulation of such regulators by Svb-REP might favor cell cycle exit and commitment towards differentiation.

Interestingly, the set of Svb target genes is markedly different in progenitors versus differentiated cells from the lineage (Figure 3F, Figure S2D), reminiscent of the situation observed between embryonic and cultured S2 cells (Figure 2). Focusing on the 195 ISC target genes, we show that their expression and regulation evolves during differentiation: 73 remain Svb targets in EB, 25 are also targets in early EC, and only 5 are targets in mature EC. Thus, the change in Svb target gene set occurs gradually (Figure 3F, Figure S2D). Accordingly, ISC target genes’ expression tends to decrease during differentiation, as well as the ability of Svb-ACT to activate their transcription (Figure 3G, Figure S2C).

It was described that ISC chromatin accessibility evolves as they differentiate into EB and EC (Aughey *et al*., 2018). We analyzed chromatin accessibility at Svb target genes’ loci in ISC, EB and mEC using CATaDa (Chromatin Accessibility Targeted DamID), a technique based on the Dam methyltransferase expression specifically in one of these cell types (Aughey *et al*., 2018). We observed that the chromatin at ISC and EB target genes is most accessible in ISC, still open in EB and closed in enterocytes (e.g. *stg* and *egfr*, Figure 3H,I). On the contrary, EC target genes tend to be more accessible in differentiated EC (*e*.*g*. Cyp9b2 Figure 3J). We tested whether this trend was global by averaging CATaDa signal from entire gene sets corresponding to cell-specific target genes (Figure 3H’,I’,L’). This confirms that, as cells differentiate, the TSSs of target genes in ISC and EB close down while those of mEC become accessible.

All together these results show that Svb target gene sets evolve gradually during intestinal cell differentiation, concomitant with changes in chromatin accessibility. This suggests that, as observed in cultured cells, the chromatin state governs Svb-accessible regions thereby constraining its transcriptional output. These results prompted us to test whether Svb isoforms whose ratio changes during differentiation (as Svb-REP accumulates), could influence chromatin organization directly, surrounding its binding sites.

### SvbREP modifies local chromatin state

To address the putative effect of Svb on chromatin landscape, we used our cultured cell system and profiled the changes in H3K27 acetylation, a key mark of active enhancers (Creyghton *et al*., 2010), in control, Svb-ACT and Svb-REP conditions. We focused on Svb-bound regions and confirmed H3K27ac enrichment at Svb peaks (Figure 4A), as previously observed with public datasets (Figure 1). Interestingly, we observed a decrease in H3K27ac in presence of Svb-REP (Figure 4A, B). The amplitude of this effect depends on the intensity of Svb peaks, as revealed by the REP/control ratio (Figure 4C), suggesting that the Svb-REP directly impinges H3K27 acetylation.

**Figure 4:**
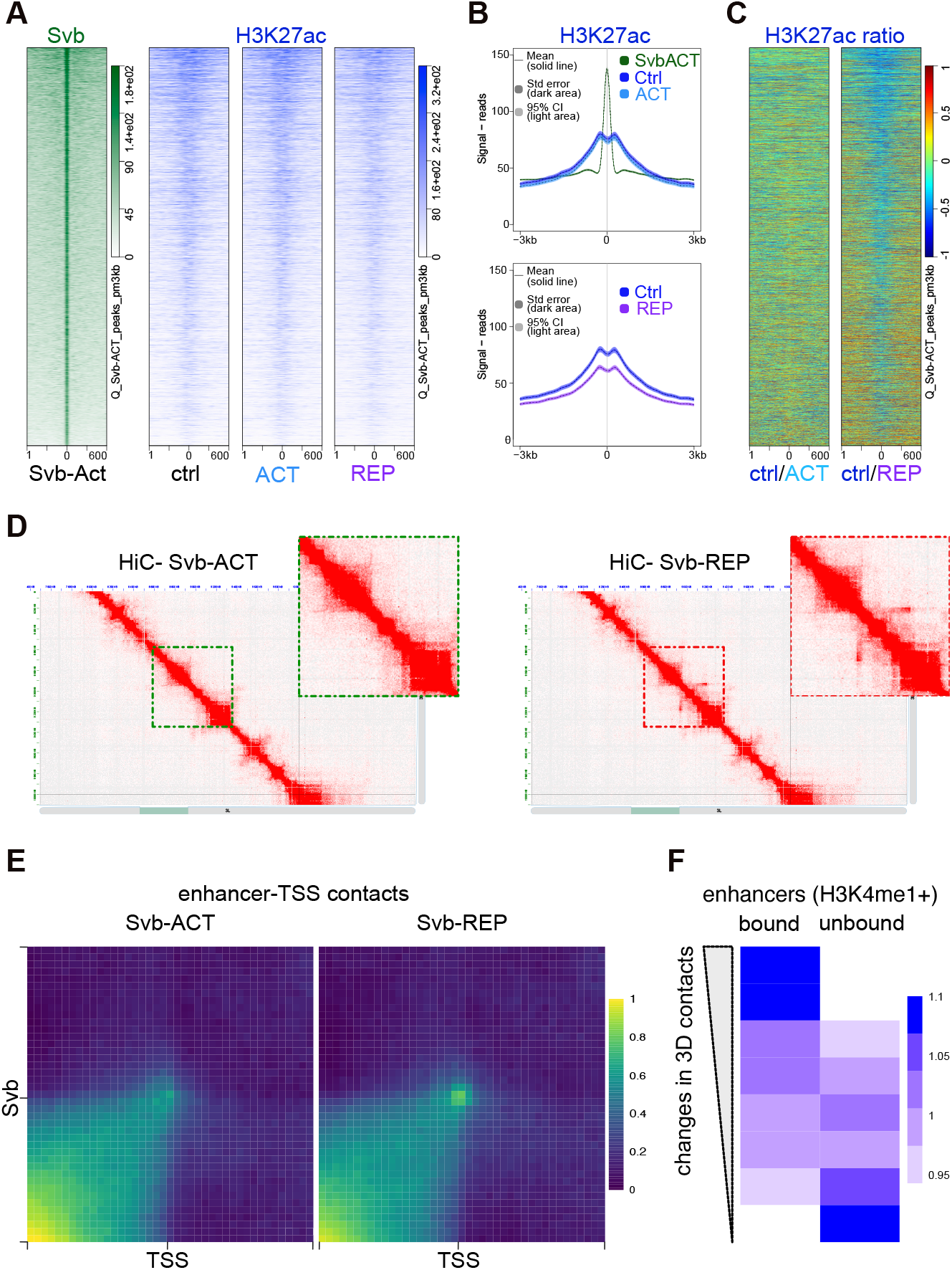
The SvbREP-to-ACT switch is accompanied by a dynamic remodeling of the enhancer chromatin landscape. (A) Heatmap showing both Svb-ACT binding (green) and H3K27ac abundance (blue) at Svb-bound regions in control, SvbACT and SvbREP cells. All heatmaps are ranked by Svb ChIP intensity. (B) Metaprofiles overlay of A (ACT vs ctrl and REP vs ctrl). (C) Heatmap showing the ratio of H3K27ac ChIP signal in ACT vs control (left) and REP vs control. (D) 2D plot showing our long-range interactions as probed using chromosome conformation capture (3C/Hi-C) in cells expressing Svb-ACT compared to Svb-REP. Enlarged areas mark possible 3D long-range contacts as highlighted (dotted squares). (E) Aggregate plots of HiC data in SvbACT and SvbREP cells assessing the dynamics of 3D chromatin contacts for all enhancers identified by STARR-seq (Arnold *et al*., 2013; Zabidi *et al*., 2015) and their distant target TSSs (see Methods). The averaged signal corresponds to the groups enriched in Svb-bound enhancers and with a decrease in long-range contacts in 3D (see panel F). (F) Genome-wide analysis of long-range contacts between enhancer and promoters in Svb-ACT compared to Svb-REP cells. Long-range contacts were measured for all putative enhancer-promoter pairs in SvbACT and compared to SvbREP cells genome-wide (see Methods). The net variations were quantilized for all enhancer-promoter pairs and ranked from highest to lowest changes in 3D contacts. A Fisher exact test was used to assess the enrichment of the quantiles of changes in 3D interactions presence or absence of Svb binding to the enhancers.

We deployed Hi-C experiments to ask whether these local changes may affect 3D chromatin organization. Inspecting long-range contacts at high resolution (1kb) using 2D plot suggested that specific 3D contacts may actually form in the Svb-REP condition, which could reflect the formation or stabilization of specific loops (Figure 4D). We wondered whether these contacts might occur between Svb bound enhancers and target promoters. To readily test this at genome-wide scales, we used aggregation plots as previously developed by us and others (Liang *et al*., 2014; Rao *et al*., 2014). Aggregate Hi-C plots depending on presence or absence of Svb suggest that, Svb-REP stabilizes such enhancer-promoter interactions (Figure 4E). To further test the specificity of these 3D interactions, we considered all enhancer-promoter interaction changes when comparing Svb-ACT and REP conditions. These couples were ranked following changes in 3D contacts. For each decile from the list, we estimated the enrichment for Svb-bound or Svb-unbound enhancers (Figure 4F and methods). This revealed that the Svb-bound enhancers tend to be enriched among the enhancers on which Svb had the strongest influence, in contrast to unbound enhancers (Figure 4F). Altogether, our data thus show that Svb-REP impinges both on local chromatin landscape of enhancers and 3D contacts with promoters. Svb-REP has an instructive role on chromatin and participates in its remodeling.

## Discussion

Here, we describe how Svb regulates the expression of its target genes. First, Svb-ACT and REP bind to the same genomic enhancer regions to antagonistically control a cohort of target genes. Second, this cohort is constrained by the chromatin environment of the cell, leading to almost completely distinct target gene sets in different cell types. Last, we show that Svb-REP isoform modifies local chromatin landscape at bound enhancers, and 3D contacts with target promoters. This work reveals how a peptide-induced proteasomal processing of a transcription factor leads to specific cell fate commitment: the crosstalk between chromatin organization and Svb activity defines cell specific transcriptional output.

In the intestine, we show that in stem cells Svb target genes (*egfr, stg*…) are activated to maintain stemness and proliferation. The progressive accumulation of Svb-REP occurring in EB will first reduce these genes’ expression, consistent with cell cycle arrest and differentiation. Notably, EGFR pathway was previously shown to directly control *svb* transcription (Al Hayek *et al*., 2021), forming a feedback loop that could reinforce the asymmetry between ISC daughter cells. In ISC, high Svb-ACT would maintain high EGFR, and high Dl, while cells starting to accumulate Svb-REP would decrease EGFR and Dl, triggering differentiation.

During differentiation, we observe that the chromatin at Svb ISC target genes’ expression goes down (Figure 3G), while chromatin around TSSs progressively closes down (Figure 3H). As we established that Svb-REP binding locally modifies H3K27ac, Svb-REP likely participates in de-acetylation of H3K27 in the intestinal lineage. This could mediate direct gene repression through inhibition of enhancer activity. In addition, this histone modification could foreshadow the chromatin compaction that occurs during differentiation, which mediates long-term silencing of “stemness” genes. It would be interesting to study the mechanisms by which Svb modulate chromatin landscape and to which extent this applies to OvoL proteins and other transcription factors.

## Acknowledgements

We thank all members of the Payre lab for critical reading of the manuscript. We thank Marion Aguirrebengoa (BigA platform) for her bioinformatic expertise, Emmanuelle Näser from the Cell sorting service of the TRI genotoul platform, Hugues Parrinello from the MGX sequencing platform. This work was supported by ANR (Chrononet, to FP’s team; Helico, OC’s team), Fondation pour la Recherche Médicale (FRM, grant DEQ20170336739 to FP’s team and DEQ20160334940 to OC’s team) and a FRBT grant. AMF, AR and DD were supported by fellowships from Ligue contre le Cancer. CI was supported by FRM and Université Paul Sabatier.

## Authors’ contributions

AMF performed ChIP-seq experiments for histone marks in S2 cells, with the help of JA, and most computational analyses with the help of AR (unsupervised clustering), DD (Histones ChIP-seq) and CI (intestinal cells RNA-seq analyses). CI and CP dissected guts of different genotypes; CI did FACS purification and prepared samples for intestinal cells RNA-seq. PL and SP prepared S2 cell samples for RNA-seq and Svb::GFP ChIP-seq. GA and TS performed CATaDa analysis. OF, NC and OC performed and analyzed HiC. OC, SP, FP and CP designed the project. FP, OC and CP supervised the project. CI, FP, OC and CP wrote the paper with contribution from all authors.

## Materials and methods

### S2 cells lines

*Drosophila* S2 cells were grown in Schneider medium supplemented with 10% fetal calf serum and 1% penicillin/streptomycin (Invitrogen). We used stable cell lines co-expressing the copper-inducible constructs pMT-Svb::GFP and pMT-pri (“1B”), or pMT-Svb::GFP and pMT-pri1-4fs (“FS”), which encodes a frame-shifted variant as a negative control (Zanet *et al*., 2015). The expression of pMT plasmids was induced by CuSO4 at the final concentration of 1 mM during 24h.

### ChIP experiments

ChIP experiments were performed essentially as previously described (Lhoumaud *et al*., 2014) with the following specifications: Cells are cross-linked with 0.8% of formaldehyde for 10 min at room temperature. Cross-linking is stopped by adding glycine (2M) and cells were washed twice in cold with PBS 1X and NaBu 1 mM. Cells are resuspended in 500 µL of ChIP permeation buffer (PBS + 0.2% Triton + 10 mM NaBu), incubated for 20 min at room temperature; cells are washed with Lysis Buffer (140 mM NaCl 5M, 15 mM HEPES pH 7.6, 1 mM EDTA, 1% Triton, 0.1% Sodium Deoxicholate, 0.5 mM DTT,10nM sodium butyrate, 25X protease inhibitor), and resuspended in Lysis Buffer, 1% SDS, 0.5% N-lauroylsarcosine. Chromatin is sonicated in Bioruptor (Diagenode) using high-power settings, intervals of 30s burst/pause for 15 cycles to obtain fragments of ≈ 250 pb. The sonicated chromatin is diluted 10 times with LB no SDS on protein low bind tubes (Eppendorf) and the chromatin fraction cleared by centrifugation (16000 g, 10 °C, 5 min) is then used for immunoprecipitation. Beads are washed and blocked with Lysis Buffer 0.1% SDS, 0.5% N-lauroylsarcosyl and BSA 0.1 mg/mL. Complex beads antibodies were established at 4° C overnight with Lysis Buffer 0.1% SDS, 0.5% N-lauroylsarcosyl on protein low bind tubes.

GFP-TRAP beads (Chromotek) and anti H3K27ac (abcam ab4729) antibodies were used for ChIP.

Chromatin is pre-clarified O/N at 4° C before the immunoprecipitation step that is carried out at 4° C for 4h. 10% of each sample is used as input. Chromatin-bead complexes are eluted twice at 70 °C during 20 min, first with 10 mM EDTA, 1% SDS, 50 mM Tris-Cl ph8 and then with TE, 0.67% SDS. Reverse cross-link is performed overnight at 65 °C and DNA purified after phenol-chloroform extraction and ethanol precipitation, for final resuspension in 1x TE.

### ChIPseq sequencing and analysis

Data of ChIPseq for H3K4me1 (GSE63518), H3K4me3, H3K27ac (GSE36374), H3K27me3 (GSE93100), H3K9me2(GSE47229), H4K16ac (GSE94115) were used to analyze the chromatin landscape of Svb binding sites. ChIP sequencing of control, FS (REP) and 1B (ACT) S2 cells (single-end 50 nt) were performed with HiSeq 2000 (Illumina) at BGI (GSE199513). Histones ChIP sequencing (paired-end 50 nt) was performed at Montpellier GenomiX platform on NovaSeq 6000 (Illumina) with NovaSeq Reagent Kits (300 cycles). The H3K27ac ChIPseq bank construction was also done by Montpellier Genomix platform with TruSeq® ChIP Sample Preparation (Illumina) (GSE199512).

For all conditions, reads were mapped on dm6 genome (r6.13 flybase) with bwa (v0.7.17-r1188) with default parameters, unique reads were filtered and peak calling was done on replicates merge files with MACS2 (Feng *et al*., 2011) for histone marks of databases (--nomodel --broad --f BAM --g dm --B --q 0.00001) or for histones mark made in this study (--nomodel --f BAMPE --g dm --B --q 1 e-05 --broad) and (-f BAM --B --g dm --to-large --nomodel --q 0.0001 or --q 0.1) for Svb forms. Metrics and quality of ChIPseq peaks were calculated with fastQC (v0.11.8) and ChIPQC package (Carroll *et al*., 2014)(v 1.26.0). To compare the set of peaks, ChIPpeakanno9 (v3.24.2) packages and BEDtools fisher (v2.29.2) were used.

ChIPseq signal was normalized by taking into account the genome mapping with bamCoverage (Ramírez *et al*., 2016) (--binSize 10 --normalizeUsing RPKM -- effectiveGenomeSize 125464728), then we normalized for the noise due to the experiment with bigwigCompare (-b1 experiment file -b2 input file --scaleFactors 1:1 --operation “log2”). Due to the fact that SvbREP is less express than SvbACT in S2 cells the normalization for the noise considers the importance of this difference scaling SvbREP peaks signal by the result of the ratio of logFC for svb in the two conditions: --scaleFactors 1.41:1. Heatmaps of binding enrichment were performed with computeMatrix (-bS 100 -a 3000 –b 3000) and plotHeatmap (deeptools v 3.5.0 (Ramírez *et al*., 2016)). RSAT (Medina-Rivera *et al*., 2015; Thomas-Chollier *et al*., 2008) was used to determine the motif enrichment on Svb peaks. Presence analysis of histones in presence of Svb were made with deeptools fisher (Ramírez *et al*., 2016) software. Coverage analysis was done with MultiBamSummary and plotCorrelation of deeptools (Ramírez *et al*., 2016) software.

### RNA Extraction, Sequencing and Analysis on S2 cells

RNAseq experiments were made on control, 1B and FS cells (GSE199511). Cells were harvested and RNA extraction was made with RNeasy Kit (Qiagen). Bank of reads and sequencing were done by HiSeq 200 (Illumina) at BGI to obtain 29-34M of reads (single-end 50nt) per replicates. The quality of sequencing was measured with fastQC (v0.11.5). Mapping was done with STAR (Dobin *et al*., 2013)(v 2.5.2b, default parameter) on Drosophila genome dm6 (r6.13 flybase), reads were counted with HTseq-count (Anders *et al*., 2015)(v0.6.0, -t gene -r pos -i gene_symbol) and statistical analysis were performed with edgeR (Robinson *et al*., 2010) using the negative binomial generalized log-linear model to the read counts for each gene (gmLRT) and decideTestDGE to determine differential genes. Graphs were obtained with Glimma (Su *et al*., 2017)(v 2.0.0) and Plotly (Sievert, n.d.)(v 4.9.3). Identification of direct target genes was performed with GenomicRanges (Lawrence *et al*., 2013)(v 1.42.0) and superExactTest (Wang *et al*., 2015). Clustering analysis described in dedicated Clustering method section.

### Drosophila genetics

See complete genotypes and references in Reagents and Tools tableTo purify ISC and EB, *esg*^*ts*^ virgin females were crossed with the following males: Control, RNAi, ACT, or REP. To purify enterocytes, *MyoIAts* virgin females were crossed to the same males. Adult progeny (only virgin female) was collected, aged at room temperature (22°c) for 72h then transferred at 29°C to induce UAS expression. Flies were kept for 6 days at 29°C, changing the food tube every 48h hours to avoid bacterial proliferation and intestinal stress.

### Gut dissection and cell dissociation

Batches of >20 virgin females were CO2-anaethesized, put on ice, their midgut were dissected in PBS. Malpighian tubules and hindgut were removed and only the posterior part of the midgut is kept (R3, R4, R5). PBS was replaced by cold 1mM PBS1X-EDTA. Posterior midguts were shrunk into pieces using a razorblade (around 30-60 seconds on a glass plate), then transferred into a fresh tube. Samples were spn 5minutes at 1000rpm 4°C, and supernatant discarded except 50µL. 200uL TrypLE 10X was added (8X final with 0.2mM EDTA) and the guts were incubated at 37°C for 10minutes with intense rocking. Digestions were stopped with 750µL SSM (Serum Supplemented Medium = Schneider’s medium +10% Fetal Bovine Serum + 1% Pen/Strep. Then samples were treated differently for FACS (ISC, EB, eEC) or for filtering (mEC). In both cases, mechanical trituration was performed by pipetting up and down using low binding, flame-rounded narrow tips (home made).

#### Sample preparation for FACS

The cell solution was strained into a cold falcon tube (70µM nylon cell strainer, pre-wet) with 1mLSSM). The Eppendorf and the strains were rinsed with 3×1mL PBS-EDTA, so that the samples are 5mL total. The quality of the dissociation and the health of the cells are checked under the macroscope. Samples were spun at 700rcf, 4°c, for >10 minutes. Supernatant was discarded except 100/200µL, to which we added 200µL PBS + 3µL clean NGS. Propidium Iodide was added to label dead cells, then cells of interest were sorted using FACS (Cell Sorter BD FACSAria Fusion in BSL3) directly into a tube containing 300µL 1X Tissue and Cell Lysis solution (MasterPure RNA purification kit) + 1µg Proteinase K [50µg/µL]. Cells were vortexed and placed on ice just after sorting.

#### Sample preparation for straining mature EC

We noted that ISC and EB were easy to dissociate but EC were not. Longer incubation in TRYPLE and harcher trituration allowed dissociation of individual EC1, but this was accompanied with high mortality. We thus reasoned that it would be less stressful for EC to simply proceed with mild dissociation of ISC and EB and simply recover patches of healthy enterocytes by simple filtering. For this procedure, digested samples are recovered on a prewet 70µM nylon cell strainer. The strained is rinsed to remove dissociated cells, then turned upside down on a petri dish and rinsed with 5mL PBS-EDTA (5mL) then 2mL SSM. Cell patches were checked under fluorescent macroscope. The samples were transferred into a tube and spun at 700 rcf at 4°C for 10 minutes. The supernatant was discarded except 150uL. This pellet was mixed with 150uL 2X Tissue and Cell Lysis solution (MasterPure RNA purification kit) + 1ug Proteinase K [50ug/uL].

### RNAseq, Sequencing and Quality Check on intestinal cells (ISC, EB, eEC and mEC)

RNAs were purified from each sample using MasterPure RNA purification kit, following manufacturer’s recommendation (including the DNAse treatment). Final RNAs were resuspended in 10uL TE and added 1uL RiboGuard (RNases inhibitor). Construction of RNA banks and sequencing was made on the Montpellier GenomiX Platform. RNA banks were done with the Ovation SoLo RNAseq System kit (Nugen). Bank validation was done with the quantification of the complementary DNA with the Standard Sensitivity NGS kit on Fragment Analyzer and with qPCR (ROCHE Light Cycler 480). The sequencing was done on NovaSeq 6000 (Illumina) with NovaSeq Reagent Kits (100cycles). Single-reads of 100nt were sequenced. ISC= GSE199510, EB=GSE220323, eEC=GSE220558, mEC=GSE220560.

The quality of sequencing was measured with fastQC (v0.11.5). Mapping was done with STAR^2^ (v 2.5.2b, default parameter) on Drosophila genome dm6 (r6.13 flybase), reads were counted with HTseq-count (Anders *et al*., 2015) (v0.6.0, -t gene -r pos -i gene_symbol) and statistical analysis were performed with edgeR (Robinson *et al*., 2010) using the negative binomial generalized log-linear model to the read counts for each gene (gmLRT) and decideTestDGE to determine differential genes. For the clustering analysis cf. Clustering method section.

### Clustering

Genes similarly expressed in all conditions (difference of reads < 50) were filtered-off. For each condition, the mean RNAseq count for the replicates was calculated, centered and reduced. We used two different algorithms for clustering using stats() package of R (Wiwie *et al*., 2015). The function hclust() for the hierarchical ascendant clustering was used with the Spearman distance and the Ward.D2 option. The function kmeans() was used for k-means clustering with default options. k-means clustering was repeated 15 times, 5 times with predefined centroids -corresponding to genes of the different clusters defined by hierarchical clustering- and 10 with random centroids. The different k-means clusters obtained by each iteration were compared with cluster_similarity (Clusteval (version 0.1)). We defined that the clustering is similar if the score equals 1.

For the S2 cells: We used for this analysis the RNAseq data from the S2 cells lines. We obtained after filtration a matrix of 3556 differential genes.

For the ISC data: We used for this analysis the RNAseq data from the ISC cells obtained by FACS method. We obtained after filtration a matrix of 817 differential genes. We determined by hierarchical ascendant clustering 5 clusters and use this parameter for the k-means clustering k = 5.

For the EB data: We used for this analysis the RNAseq data from the EB cells obtained by FACS method. We obtained after filtration a matrix of 1461 differential genes. We determined by hierarchical ascendant clustering 5 clusters and use this parameter for the k-means clustering k = 5.

For the eEC data: We used for this analysis the RNAseq data from the eEC cells obtained by FACS method. We obtained after filtration a matrix of 1694 differential genes. We determined by hierarchical ascendant clustering 5 clusters and use this parameter for the k-means clustering k = 5.

For the mEC data: We obtained after filtration a matrix of 1186 differential genes. We determined by hierarchical ascendant clustering 4 clusters and use this parameter for the k-means clustering k =4.

To determine the most robust group of genes we define a score of robustness or r-score. This r-score is calculated for each comparison between clusters of k-means and hierarchical algorithms. First, we define a percent of similarity between k-means clustering and hierarchical clustering such as*:((number of common gene)/((number of gene in kmeans cluster+number of gene in hirarchical cluster)-number of common gene))×100*

Then for each k-means, if this percentage is greater than or equal to the average obtained with all the clustering groups (in percentage), the comparison obtained a score of 1. For a cluster, all of these scores were added together to obtain the match score. For each cluster, we also calculated the intersection, conservation and dispersion. intersection (i)= minimal number of genes common to all clustering (h-clustering and all k-means). conservation (c)= i/ number of genes in the h-clustering. dispersion (d)= higher number of genes corresponding to one cluster/i. Finally the r-score = c/d * match score

### Catada Analysis

CATaDa chromatin accessibility data from ISCs, EBs, and ECs were obtained from (Aughey *et al*., 2018). Per-GATC fragment counts were RPM normalised and visualised in IGV. GFF signal files were converted to bigwig format using kentUtils bedGraphToBigWig and profiles of RPM chromatin accessibility scores across transcriptional start sites were plotted using deeptools computeMatrix and plotProfile functions. Chromatin accessibility at individual promoter regions for global comparison were extracted using 2kb regions centered on the TSS, means were compared using Kruskal-wallis test with Dunn’s multiple comparison.

### HiC Analysis

Hi-C analysis at high resolution (1kb) was performed for cells expressing either Svb-REP or Svb-ACT (GSE221863) and performed as previously described (Heurteau *et al*., 2020) using pipelines available through Github https://github.com/CuvierLab/Hi-C data for analysis of 3D genomic interactions in Drosophila S2 cells. Aggregation analysis was performed as previously in 1D/2D plots (Liang et al., 2014; Rao et al., 2014) to compare genomic contacts in cells expressing either Svb-REP or Svb-ACT. Hi-C data were aggregated onto pairs of enhancers identified by STARR-seq (Arnold *et al*., 2013; Zabidi *et al*., 2015) and distant target transcription start sites. Enhancer-promoter pairs were then ranked depending on net variations of 3D contacts between Svb-REP and Svb-ACT and the ranked quantiles of pairs were assessed for enrichment depending on presence or absence of Svb binding or not using a Fisher exact test.

