## Supplementary Figures for "Crosstalk between chromatin and the transcription factor Shavenbaby defines transcriptional output along the *Drosophila* intestinal stem cell lineage"

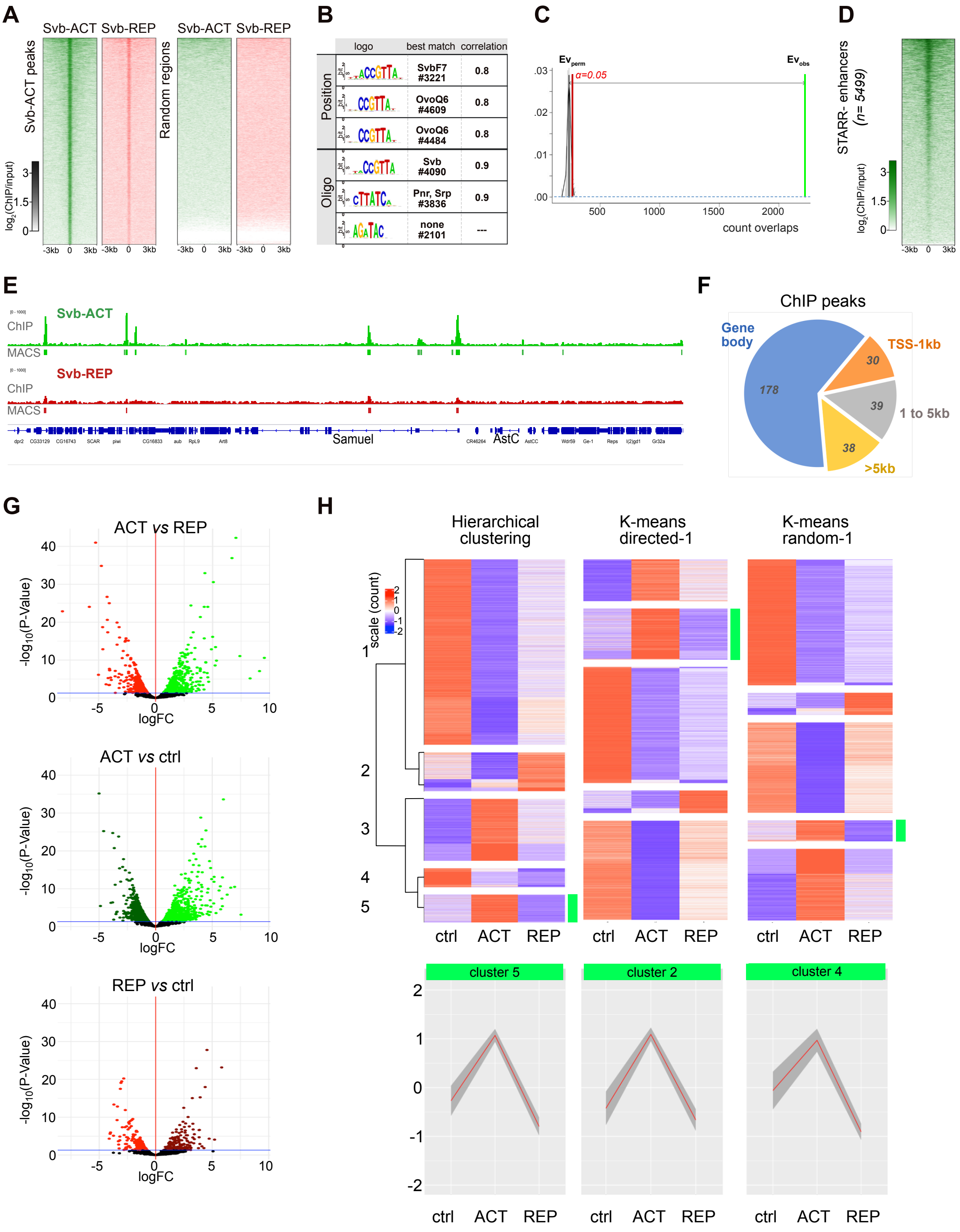


**Figure Supp 1**

**Figure S1: Mode of action of Svb-ACT and REP to bind DNA and regulate their target genes**

(A) ChIP-seq heatmaps for Svb-ACT (green) and Svb-REP (red) enrichment across all significant peaks of the *Drosophila* genome detected in Svb-ACT cells (n=5512). Each row represents a 6 kb window centered on peak midpoint, sorted by Svb-ACT ChIP-seq signal. ChIP-seq signal of a similar set of random regions is show on the right. All heatmaps are centered on Svb-ACT peaks and sorted by Svb-ACT signal intensity. Color intensity reflects the level of ChIP-seq enrichment presented as log2(ChIP/input).

(B) DNA motifs enriched in Svb-ACT bound peaks, as defined by two algorisims (position or oligo) of the RSAT suite (Nguyen *et al.*, 2018).

(C) Hypergeometric test (Z score =129.05, p-value= 0.0099) on the data set from figure 1E (grren) compared to 100 random permutations (grey) of the data. X axis represents the number of overlaps, y axis the observed frequency.

(D) The heatmap shows the enrichment of Svb-ACT at STARR-S2 enhancer regions (5,499); each row represents a 6kb window centered on STAR-seq peak midpoint, sorted by the intensity of Svb-ACT ChIP signal (Arnold *et al.*, 2013).

(E) Gene browser screenshot of the *Samuel* locus, showing respective intensity of Svb-ACT and Svb-REP signals.

(F) Location of Svb peaks within the set of 285 target genes obtained with the supervised method. Gene body represents the whole transcribed region extended from 300 pb upstream the TSS down to + 1kb downstream the transcriptional stop; TSS-1kb, to 1kb region upstream of the TSS, 1 up 5kb, the region 1-5kb upstream of TSS. Other peaks are located at distances >5kb upstream of TSS.

(G) Volcano plots show differentially expressed genes (DE) upon Svb-ACT or Svb-REP expression in S2 cells. Control (ctrl) condition is S2 parental cells.

(H) Heatmaps represent cluster of DE genes in S2 cells, using hierarchical clustering and directed or random k-means unsupervised appoches, based on mRNA expression levels in control (ctrl), Svb-ACT and Svb-REP conditions. Each horizontal line represents a gene averaged from two replicates in each condition. Expression level (count) is color coded. Clusters comprising Svb target genes are highlighted by green boxes. The charts (meta-profiles) below represent cumulative expression of all genes of the cluster in the three conditions.

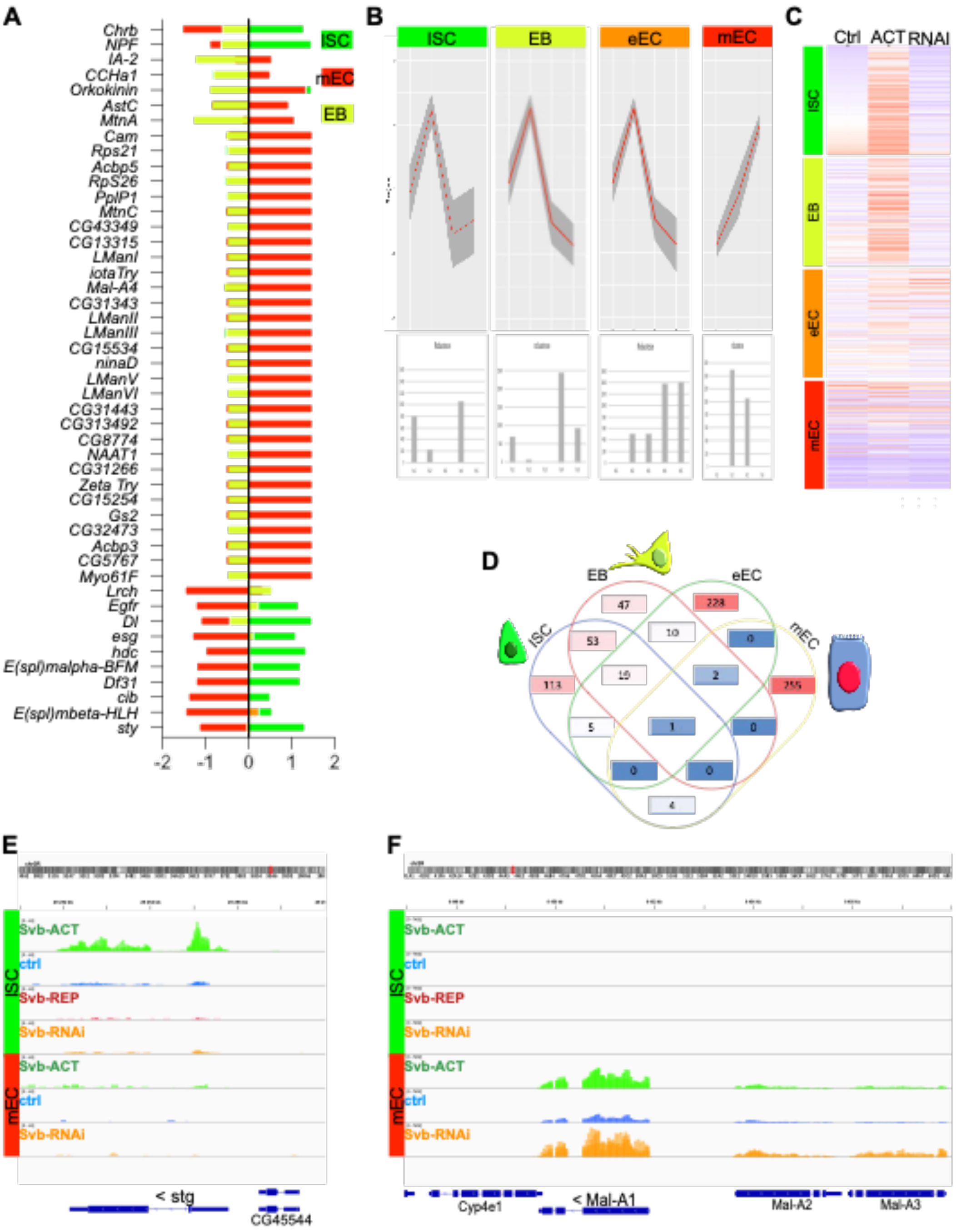


**Figure Supp 2**

**Figure S2: Identification of Svb target genes in the intestinal lineage and analysis of their behavior.**
(A) Graph showing the behavior of the top 10 genes as defined in the cell atlas (ref., set 2) for the precursors (ISC+EB) and the different differentiated ECs located in the hind midgut, within the RNA-seq corresponding to the different cell types.
(B) Top, metaprofiles of the cluster corresponding to Svb target genes for each intestinal cell types. Bottom, graph showing the robustness (r-score) of each cluster in the corresponding cell type.
(C) Heat map showing the evolution of the expression of Svb target genes during the differentiation of ISCs into ECs.
(D) Venn diagram analysis the intersection of the set of Svb target genes identified in each cell type.
IGV snapshots of the *stg* (E), *Mal-A1*(F) loci are shown (representative of ISC and mEC target genes, respectively).
